# Debutylation and Dephenylation of Organotin Compounds Through Fungal Laccase–Mediator Systems

**DOI:** 10.1101/2025.09.25.678422

**Authors:** Chin-Feng Chang, Chuan-Hsin Ho, Jun-You Liu, Shiu-Mei Liu

**Affiliations:** Department of Biological Science and Technology, China University of Science and Technology, Taipei 11581, Taiwan; Department of Science Education, National Taipei University of Education, Taipei 10671, Taiwan; Institute of Marine Biology, National Taiwan Ocean University, Keelung 20224, Taiwan

**Keywords:** Tributyltin, triphenyltin, laccase, mediator, white rot fungi

## Abstract

In this study, the biodegradation of tetrabutyltin (TeBT) and tetraphenyltin (TePhT) through various laccase–mediator systems were investigated. The fungal laccases used were from the extracellular fluid of solid fermentation culture of white-rot fungi *Perenniporia tephropora* TFRI 707, *Agrocybe aegerita*, and *Pleurotus geesteranus*, with 2,2’-azino-bis (3-ethylbenzthiazoline-6-sulfonic acid), violuric acid, 4-hydroxyacetophenone, and syringaldehyde being used as mediators. After 7 days of incubation, 300 μg/L of TeBT or 1000 μg/L of TePhT were added, the laccase obtained from *P. tephropora* TFRI 707 removed 83.9%–87.4% of TeBT and 78.0%–91.9% of TePhT; the laccase obtained from *A. aegerita* removed 73.7%–88.9% of TeBT and 75.5%–92.0% of TePhT; and the laccase obtained from *Pl. geesteranus* removed 70.4%–85.3% of TeBT and 72.9%–90.2% of TePhT (percentages include incubation with and without mediators). Tributyltin (TBT), dibutyltin (DBT), and monobutyltin (MBT) were detected during the degradation of TeBT. Triphenyltin (TPhT), diphenyltin (DPhT), and monophenyltin (MPhT) were detected during the degradation of TePhT. Therefore, laccases can effectively and rapidly remove butyl or phenyl side chains from the organotin compounds. Because dealkylated alkyl organotin compounds and the elemental tin exhibit considerably low toxicity, laccases can potentially be used in environmental cleanup.

## INTRODUCTION

Organotin compounds, such as tributyltin (TBT) and triphenyltin (TPhT), have harmful effects on various organisms, even at low nanomolar aqueous concentrations (1). They are also suspected to have an endocrine-disrupting effect in humans (2–4). Organotin compounds can be removed through abiotic and biotic degradation. The abiotic degradation of organotin compounds, which involves processes such as photolysis, chemical cleavage, and thermal cleavage, has a negligible effect on the natural breakdown of TBT (5). Considerable evidence suggests that biotic degradation is the major pathway for the removal of organotin compounds in aquatic and sediment environments (6). Microorganisms such as microalgae, fungi, and bacteria are responsible for the biotic degradation of TBT and TPhT (7–10). Furthermore, TBT and TPhT can be biodegraded through the progressive removal of the alkyl side chains from a tin atom, which results in the successive production of di-substituted (R_2_SnX_2_), mono-substituted (RSnX_3_) compounds, and inorganic tin (11,12). In a previous study, tri-substituted TBT (R_3_SnX) organotin was the most potent substrate for industry application, and the toxicity of the members of the butyltin family decreased in the following order: TBT > DBT > MBT > tin (6). The toxicity of phenyltin compounds has been considered the same as for butyltins, that is TPhT > DPhT > MPhT > tin (8). However, only a limited number of microorganisms possessing a TBT degradation ability or TPhT degradation ability have been identified (7).

Fungal laccases, especially those produced by white-rot fungi, have the potential to be used in the remediation of toxic compounds (13–16). These laccases belong to a family of enzymes known as multicopper oxidases and can oxidize various compounds (17). However, their application was previously known limited to certain xenobiotic compounds because of their low oxidation potential (18). In the presence of mediator compounds, such enzymes, fungal laccases exhibit a relatively high oxidation capability, which enables these laccases to oxidize a wider variety of xenobiotic compounds. Fungal laccase–mediator systems (LMSs) have been applied in pulp delignification (19,20), textile dye bleaching (21), polycyclic aromatic hydrocarbon degradation (22,23), pesticide or insecticide degradation (24,25), organic synthesis (26), and the bioremediation and detoxification of aromatic pollutants (27,28).

In this study, the biodegradation of butyltin and phenyltin compounds through various LMSs was investigated. The fungal laccases used were produced from *Perenniporia tephropora* TFRI 707, *Agrocybe aegerita*, and *Pleurotus geesteranus*, with 2,2’-azino-bis(3-ethylbenzthiazoline-6-sulfonic acid) (ABTS), violuric acid (VLA), 4-hydroxyacetophenone (4-HAP), and syringaldehyde (SIR) being used as mediators. This study investigated the potential of fungal LMSs to degrade butyltin and phenyltin compounds and the necessity of using a mediator.

## MATERIALS AND METHODS

### Chemicals

Crude laccases originate from the solid fermentation culture fluid of the white-rot fungi *P. tephropora* TFRI 707, *A. aegerita*, and *Pl. geesteranus*. TBT, DBT, and MBT were obtained from Sigma-Aldrich (St. Louis, USA), tetrabutyltin (TeBT) was obtained from Acros Organics (Geel, Belgium). Stock solutions of butyltin compounds were prepared through dissolution in methanol. TPhT, diphenyltin (DPhT), and monophenyltin (MPhT) were obtained from Aldrich Chemical Co. (Milwaukee, USA). Tetraphenyltin (TePhT) was obtained from Fluka Chemicals (Vienna, Austria). Stock solutions of phenyltin compounds were prepared through dissolution in ethanol. Sodium tetraethylborate (NaBEt_4_, 98%) was obtained from Strem Chemicals (Bischheim, France). SIR was obtained from Acros Organics. ABTS (98%) was obtained from Sigma-Aldrich; VLA (97%) was obtained from Fluka Chemicals; and 4-HAP was obtained from Acros Organics. All the other chemicals and solvents were high-performance liquid chromatography or reagent grade. A manual solid phase microextraction device was obtained from Supelco (Bellefonte, USA).

### Incubation of white-rot fungi and the collection of crude enzymes

The fungal strain *P. tephropora* TFRI 707, *A. aegerita*, and *Pl. geesteranus* were maintained in potato dextrose agar (PDA; Difco, Detroit, USA) at 4 °C. Agar plugs cut from the outer circumference of an actively growing fungal colony on PDAs were used as inocula. Solid fungi incubation was performed in 2800 mL Fernbach flasks. Each flask containing 320 mL of culture medium was inoculated with 40 agar plugs (diameter of 7 mm) and incubated without shaking at 30 °C. For the fungal strain *P. tephropora* TFRI 707 the culture medium (1 L) contained 25% (w/v) milled bark of the tapa tree *Broussonetia papyrifera*, 0.05 g of Tween 80, 10 g of glucose, 10 mL of 35 μM CuSO_4_ · 5H_2_O, 1 mL of trace-element solution, and 50 mM sodium tartrate buffer (pH 6.0). The trace-element solution (1 L) contained 0.08 g of CuSO_4_ · 5H_2_O, 0.05 g of FeSO_4_ · 7H_2_O, 4.3 g of ZnSO_4_ · 7H_2_O, 0.07 g of MnSO_4_ · H_2_O, and 0.076 g of Na_2_MoO_4_ · 2H_2_O. Extracellular fluid was collected every alternate day over 5 to 30 days to check laccase enzyme activities. About 14-17 days when maximum enzyme production occurred, the crude laccase enzyme solution was collected and diluted with 50 mM sodium tartrate buffer (pH 5.0). The crude laccase enzyme solution was separated from mycelia by filtration on Whatman paper.

The laccase enzymes of *A. aegerita* and *Pl. geesteranus* were extracted from the spent mushroom substrate with the same volume of the culture medium, after 70% sawdust and 30% rice bran of bulk bag cultivation.

### Laccase assay

Laccase activity was assayed by monitoring the oxidation of ABTS in a reaction mixture solution (15). Measurements were made in a reaction mixture solution containing 10 µL of crude enzyme sample, 490 µL of 102 mM sodium tartrate buffer (pH 3.0) and 500 µL of 2 mM ABTS. The oxidation of ABTS was monitored at 420 nm, and enzyme activity was expressed in international units (U). One unit activity of a laccase was defined as 1 µM of ABTS oxidized per minute (□ _420_ = 36000 M^−1^ cm^−1^) (29).

### Degradation rates of TeBT and TePhT when using the enzyme solution

Three sources of laccase enzymes, namely *P. tephropora* TFRI 707, *A. aegerita*, and *Pl. geesteranus*, and four mediators, namely ABTS, VLA, 4-HAP, and SIR, were used in the present study. The control blank included 40 mM sodium tartrate buffer (pH 4.0), 20% glycerol, and 300 μg/L of TeBT or 1000 μg/L of TePhT; however, no enzymes or mediators were added. The experimental group was divided into two groups, one group with mediator addition and one group without mediator addition. Moreover, 1 mM mediator and 800 U/mL of laccases were used in the experiment. All the reactions were performed in triplicate, and the reaction mixture was incubated in a sealed bottle in the dark at 25 °C under a rotation speed of 160 rpm.

### Analytical methods

Headspace solid-phase microextraction and gas chromatography (GC)-flame photometric detection (FPD) were used for the quantitative determination of the butyltin (TeBT, TBT, DBT, and MBT) and phenyltin (TePhT, TPhT, DPhT, and MPhT) compounds described in previous studies (30,31). The test sample was incubated in a sealed brown bottle at 25 °C for 50 min to allow for in-situ NaBEt_4_ derivatization and extraction to the needle of a microextraction fiber. The needle was immediately inserted into a GC injector for thermal desorption. Quantitative determination of butyltin and phenyltin compounds was performed using a Dani 1000 GC equipped with a column (HP–5, 30 m × 0.25 mm i.d. × 0.25 μM film thickness, Hewlett Packard, USA) and a FPD fitted with a 610-nm optical filter. The fiber was injected in splitless mode. The column temperature was programmed to increase from 70 °C (holding time of 1 min) to 190 °C at a rate of 30 °C/min, 270 °C (holding time of 3 min) at a rate of 15 °C/min, and 290 °C (holding time of 1 min) at a rate of 15 °C/min. The injector temperature was 250 °C, and the detector temperature was 290 °C. All the experiments were performed in triplicate.

## RESULTS

### Laccase activity profiles during the solid fermentation of the fungal strain *P. tephropora* TFRI 707

The activity of the laccase obtained from the extracellular fluid of *P. tephropora* TFRI 707 was 4352.9 U/mL after 5 days incubation, and it has the highest enzyme activity 6025.7 U/mL by the 17th day (Fig. 1A). The maximum protein concentration of this laccase was 1.22 mg/mL (Fig. 1B). The enzyme activities of the laccases obtained from *A. aegerita* and *Pl. geesteranus* samples were 1748.6 and 811.4 U/mL, respectively.

**Fig. 1.**
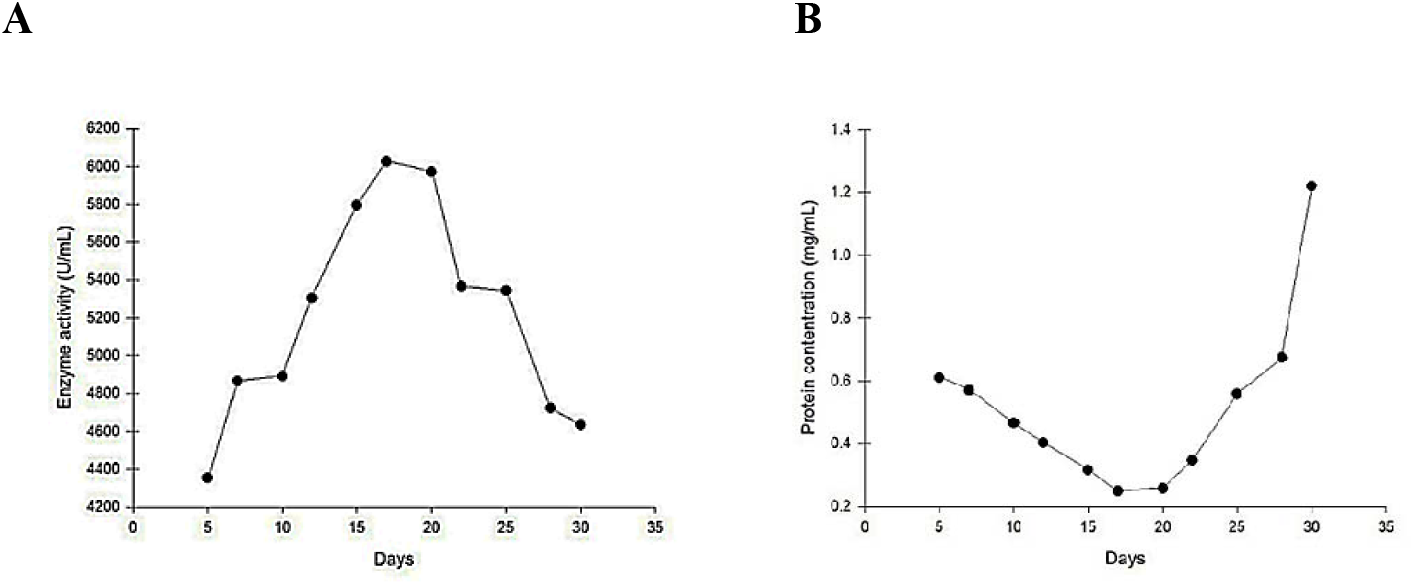
Laccase activity (A) and protein concentration (B) in the extracellular fluid during solid fermentation of *P. tephropora* TFRI 707.

### Degradation of TeBT by the laccases obtained from *P. tephropora* TFRI 707

No degradation trend of TeBT was observed from 0 to 168 h of incubation, and no degradation products, such as TBT, DBT, or MBT, were detected in the blank batches that did not contain a laccase or a mediator. Only a 2% descending trend for TeBT was detected at 168 h in the TeBT blank batches (Fig. 2A). However, TeBT was degraded by a laccase in the test groups without mediator addition (Fig. 3A1). Only TeBT (279.39 ± 1.63 μg/L) was detected at the beginning. After 24 h, the concentration of TeBT decreased to 69.46 ± 3.69 μg/L, and TBT (5.02 ± 0.27 μg/L), DBT (4.40 ± 0.13 μg/L), and MBT (31.25 ± 1.72 μg/L) were detected. After 72 h, the concentration of TeBT decreased to 50.30 ± 4.14 μg/L, that of TBT decreased to 3.18 ± 0.27 μg/L, and those of DBT (15.76 ± 0.32 μg/L) and MBT (45.19 ± 2.02 μg/L) gradually increased. At 168 h, the concentrations of TeBT (44.93 ± 2.32 μg/L) and TBT (0.9 7 ± 0.07 μg/L) decreased to their lowest values, whereas those of DBT (17.77 ± 0.38 μg/L) and MBT (59.13 ± 1.58 μg/L) reached their highest values. The concentration of TeBT decreased by approximately 83.9% after 168 h (Fig. 3A1).

**Fig. 2.**
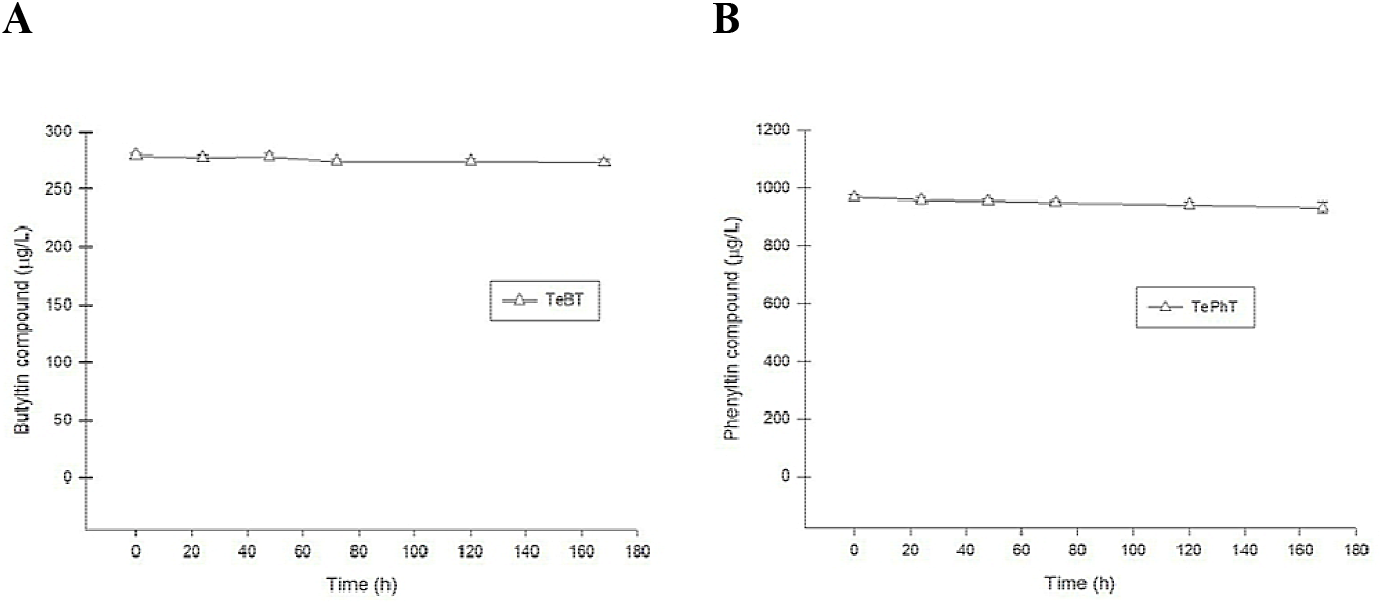
Degradation experiments of the TeBT (A) and TePhT (B) without adding laccase and a mediator. TeBT (tetrabutyltin), TePhT (tetraphenyltin).

**Fig. 3.**
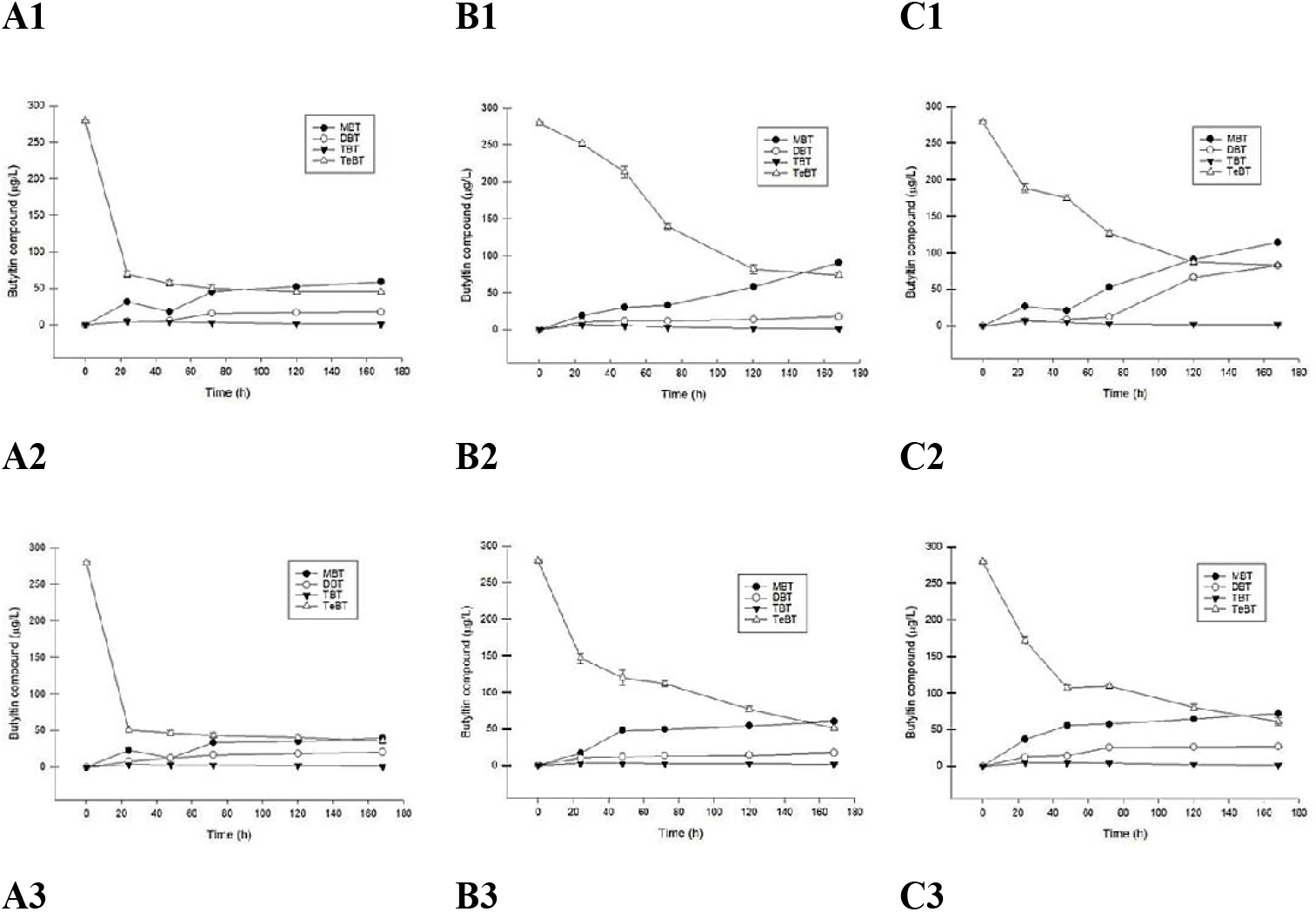
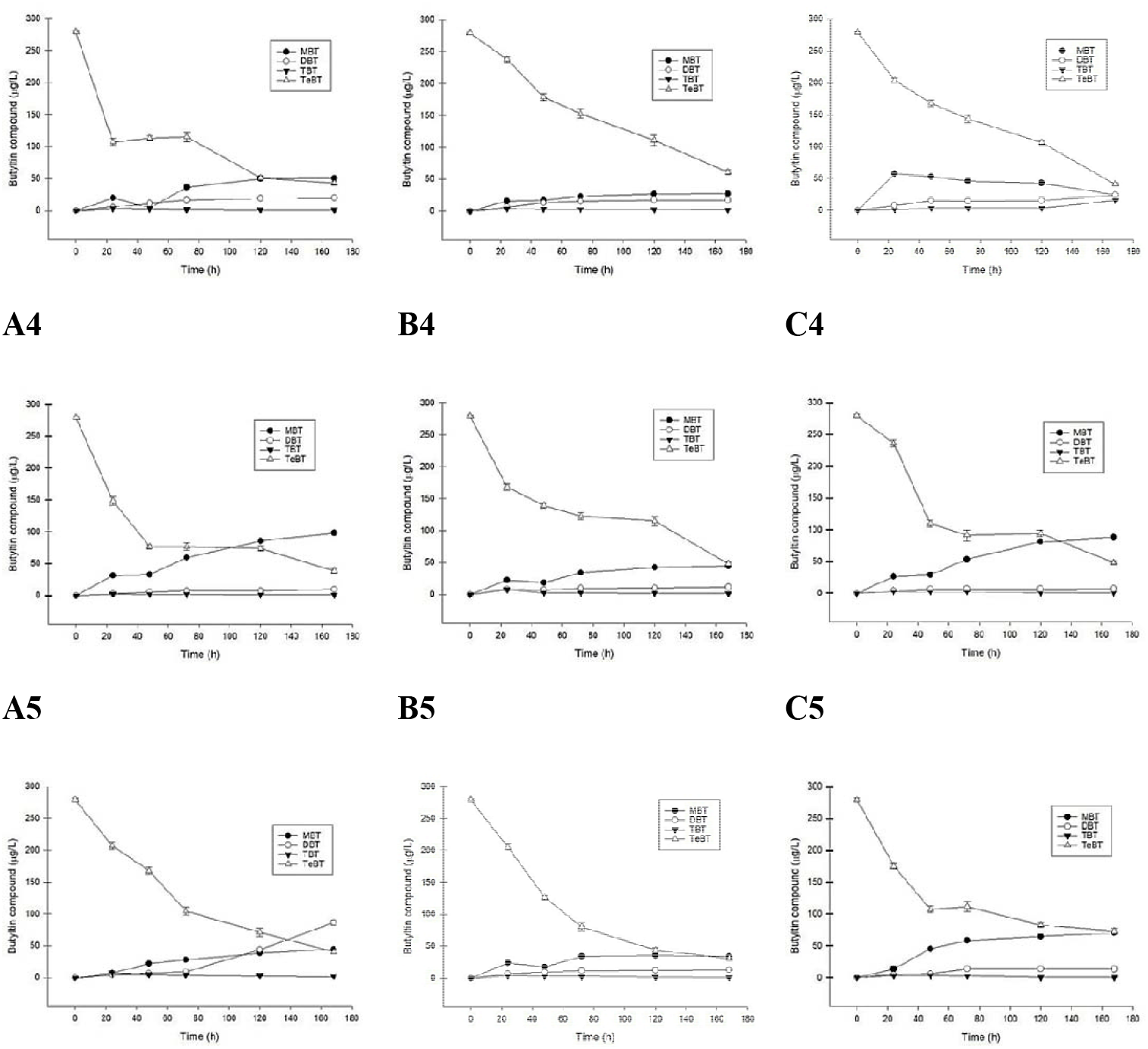
Degradation experiments of the TeBT by (A) *P. tephropora* TFRI 707, (B) *A. aegerita*, and (C) *Pl. geesteranus* laccase with/without adding a mediator during 7 days of incubation in the synthetic medium. (1) laccase alone, (2) Laccase and mediator ABTS, (3) Laccase and mediator VLA, (4) Laccase and mediator 4-HAP, (5) Laccase and mediator SIR. MBT (monobutyltin), DBT (dibutyltin), TBT (tributyltin), TeBT (tetrabutyltin).

In the groups to which a laccase and mediators (ABTS, VLA, 4-HAP, or SIR) were added, during 168 h of incubation, the concentrations of TeBT gradually decreased to 50.42 ± 2.56, 107.28 ± 5.53, 148.29 ± 6.72, and 207.07 ± 5.78 μg/L, respectively. TBT, DBT, and MBT were detected with the following concentrations: TBT, 3.473 ± 0.195, 3.201 ± 0.091, 2.894 ± 0.180, and 7.005 ± 0.296 μg/L; DBT, 7.288 ± 0.106, 6.087 ± 0.324, 2.716 ± 0.056, and 4.973 ± 0.100 μg/L; and MBT, 22.903 ± 0.690, 19.669 ± 0.756, 31.260 ± 1.911, and 6.558 ± 0.499 μg/L. After 48 h, in the mediator batches of ABTS, 4-HAP, and SIR, the concentrations of TeBT and TBT were still gradually decreasing. At 168 h, the concentrations of TeBT and TBT decreased to their lowest levels, whereas those of DBT and MBT reached their highest levels. In the VLA mediator batches, no TeBT degradation trend was observed from 24 to 72 h. Subsequently, the concentrations of TeBT and TBT gradually decreased. At 168 h, the concentrations of TeBT and TBT decreased to their lowest levels, whereas those of DBT and MBT decreased to their highest levels. The TeBT content decreased by approximately 87.4%, 84.7%, 86.3%, and 85.5% after 168 h in the mediator batches of ABTS, 4-HAP, VLA, and SIR, respectively (Fig. 3A2–3A5).

### Degradation of TeBT by the laccases obtained from *A. aegerita*

After 24 h, the incubation concentration of TeBT in the test samples gradually decreased, and TBT, DBT, and MBT were detected. Subsequently, the concentrations of TeBT and TBT continued to gradually decrease, and the concentrations of DBT and MBT gradually increased. At 168 h, the concentrations of TeBT and TBT decreased to their lowest levels, whereas those of DBT and MBT reached their highest levels. The TeBT concentration decreased by approximately 73.7% in the experimental batches of laccases without a mediator and by approximately 81.7%, 78.2%, 82.8%, and 88.9% in the experimental batches of laccases with the ABTS, 4-HAP, VLA, and SIR mediators, respectively, after 168 h (Fig. 3B1–3B5).

### Degradation of TeBT by the laccases obtained from *Pl. geesteranus*

After 24 h, the concentration of TeBT gradually decreased, and TBT, DBT, and MBT were detected. Subsequently, the concentrations of TeBT and TBT continued to gradually decrease, and the concentrations of DBT and MBT gradually increased. At 168 h, the concentrations of TeBT and TBT decreased to their lowest levels, whereas those of DBT and MBT reached their highest levels. The TeBT concentration decreased by approximately 70.4% in the experimental batches of laccases without a mediator and by approximately 78.3%, 85.3%, 82.8%, and 73.9% in the experimental batches of laccases with the ABTS, 4-HAP, VLA, and SIR mediators, respectively, after 168 h (Fig. 3C1–3C5).

### Degradation of TePhT by laccases obtained from *P. tephropora* TFRI 707

No degradation trend of TePhT was observed from 0 to 168 h in the blank batches without laccases or a mediator, and no degradation products of TPhT, DPhT, or MPhT were detected. Only a 3.6% descending trend for TePhT was observed at 168 h in the TePhT blank batches (Fig. 2B). In the test groups to which the laccases obtained from *P. tephropora* TFRI 707 were added with or without a mediator (Fig. 4A1–4A5), only TePhT was detected at the beginning. After 24 h, the concentration of TePhT decreased, and TPhT, DPhT, and MPhT were detected. Subsequently, the concentrations of TePhT and TPhT continued to gradually decrease, and the concentrations of DPhT and MPhT gradually increased. After 168 h, the TePhT concentration decreased by approximately 78.0% in the experimental batches of laccases without a mediator and by approximately 91.5%, 91.9%, 88.4%, and 87.7% in the experimental batches of laccases with the ABTS, 4-HAP, VLA, and SIR mediators, respectively (Fig. 4A1–4A5).

**Fig. 4.**
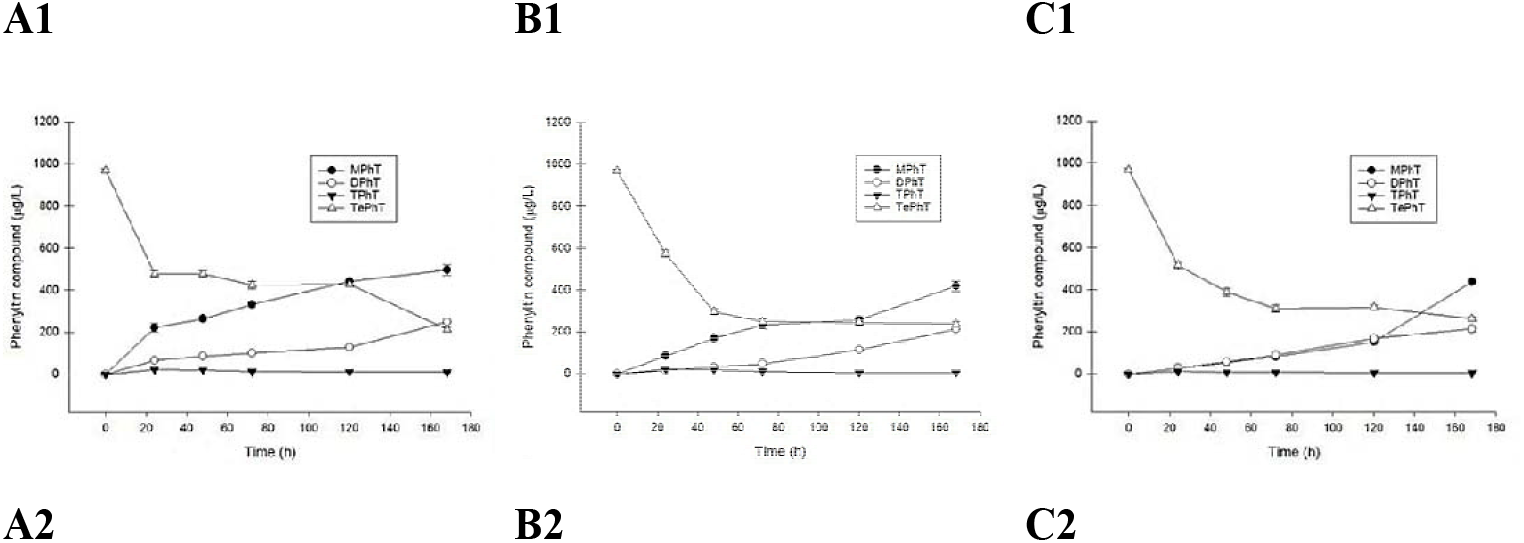
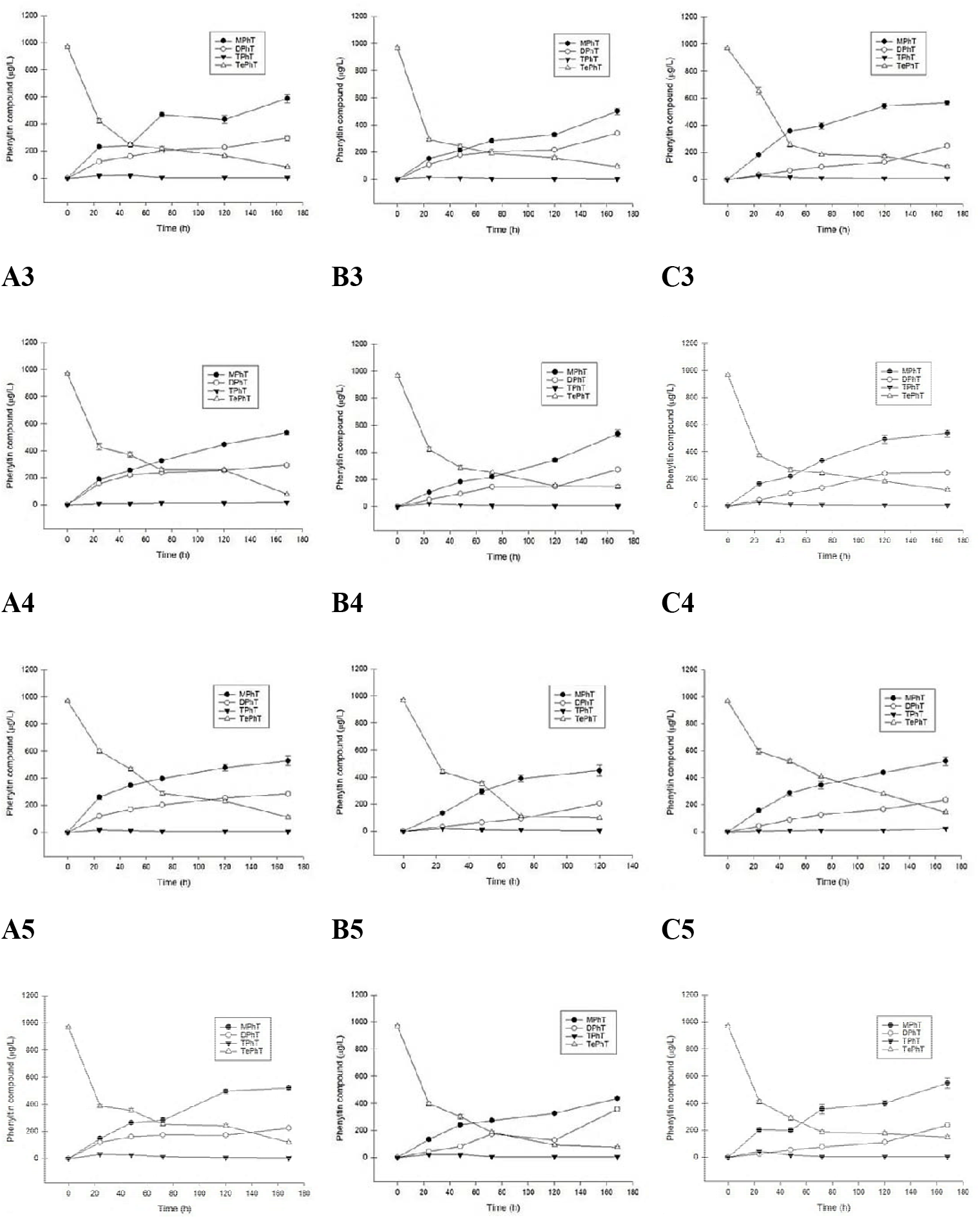
Degradation experiment of the TePhT by (A) *P. tephropora* TFRI 707, (B) *A. aegerita*, and (C) *Pl. geesteranus* laccase with/without adding mediator during 7 days of incubation in synthetic medium. (1) laccase alone, (2) Laccase and mediator ABTS, (3) Laccase and mediator VLA, (4) Laccase and mediator 4-HAP, (5) Laccase and mediator SIR. MPhT (monophenyltin), DPhT (diphenyltin), TPhT (triphenyltin), TePhT (tetraphenyltin).

### Degradation of TePhT by the laccases obtained from *A. aegerita*

In the test groups to which the laccases obtained from *A. aegerita* were added with or without a mediator (Fig. 4B1–4B5), a similar trend was observed: only TePhT was detected at the beginning. After 24 h, the concentration of TeBT gradually decreased, and TPhT, DPhT, and MPhT were detected. After 168 h, the TePhT concentration decreased by approximately 75.5% in the experimental batches of laccases without a mediator and by approximately 90.3%, 84.8%, 92.0%%, and 92.0% in the experimental batches of laccases with the ABTS, 4-HAP, VLA, and SIR mediators, respectively (Fig. 4B1–4B5).

### Degradation of TePhT by the laccases obtained from *Pl. geesteranus*

In the groups in which TePhT was degraded by laccases obtained from *Pl. geesteranus* with or without a mediator (Fig. 4C1–4C5), a similar trend was observed: only TePhT was detected at the beginning. After 24 h, the concentration of TePhT gradually decreased, and TPhT, DPhT, and MPhT were detected. After 168 h, the TePhT concentration decreased by approximately 72.9% in the experimental batches of laccases without a mediator and by approximately 90.2%, 87.8%, 85.2%, and 84.7% in the experimental batches of laccases with the ABTS, 4-HAP, VLA, and SIR mediators, respectively (Fig. 4C1–4C5).

## DISCUSSION

Several studies have attempted to identify the effects of microorganisms on TPhT bio-removal (8, 32–35). Although biodegradation of TPhT has been demonstrated by these studies, information on the microorganisms involved in biodegradation is still considerably limited. In a 231-day incubation experiment, TPhT degradation was found to be considerably slower in sterile soil than in a nonsterilized sample (8). Liu et al. (33) reported that DBT, MBT, and inorganic tin (Sn) were produced simultaneously following TBT degradation in estuarine sediment slurries. The highest TBT degradation rates in sediment slurries (15–38 μg/L/day) were considerably higher than those in aseptic control sediment slurries (2.6–3.8 μg/L/day). Therefore, the biotic process was the most crucial mechanism for TPhT and TBT degradation in soil and estuarine sediment slurries, respectively. Shuto et al. (32) screened microbial strains capable of degrading TPhT and isolated *Pseudomonas* sp. that could tolerate TPhT at 50000 μg/mL. In a synthetic medium containing 1000 μg/mL of TPhT and 0.16% ethanol, the strain degraded TPhT by a maximum of 50% over 5 days and produced the following by-products: 38% DPhT, 12% MPhT, and 1.5% inorganic tin. However, the degradation of TPhT did not occur in the nutrient medium; therefore, the authors suggested that TPhT might be cometabolized with ethanol. In a previous study, the removal efficiency of 500 μg/L of TPhT after its degradation by a 0.3 g/L biomass of the bacterium *Brevibacillus brevis* for 5 days was approximately 60%; however, suitable levels of oxygen as well as suitable types and levels of nutrients, surfactants, and metals improved TPhT degradation by 15%–25% (34). Gao et al. (35) reported the biosorption, degradation, and removal efficiencies of TPhT (0.5 mg/L) by 0.3 g/L viable cells of the bacterium *Stenotrophomonas maltophilia* at 10 days to be 3.8%, 77.8%, and 86.2%, respectively, and the adsorption efficiency of inactivated cells was 72.6%. Most studies on the degradation of organotin compounds have used bacteria, and few studies have directly used enzymes. Strain LBA6, an *Achromobacter* sp. isolated from TBT-contaminated sediment, is the first record of producing enzymes that degrade 25% of the initial TBT concentration (36). The laccases used in the present study can degrade organotin compounds at room temperature, and 45.7 U/mL of enzyme activity remained after 1 week, which indicated that the laccases have a high tolerance to environmental factors.

Applications of laccases alone have frequently been limited to the oxidation of recalcitrant substrates with high redox potential (19). One report mentioned that the treatment of the antibiotics sulfadimethoxine (SDM) and sulfamonomethoxine (SMM) with the laccases obtained from *Perenniporia* sp. TFRI 707 alone resulted in the limited reduction of these antibiotics. The ABTS, VLA, and SIR mediators were effective in the oxidation of SDM and SMM in the LMSs. Under optimal conditions, the *t*_1/2_s values of the two sulfonamide antibiotics were demonstrated to be in the range of 0.4–2.4 min in LMSs containing ABTS, VLA, and SIR (28). Therefore, several redox mediators were added in this study with laccases to form LMSs to expand the application potential of laccases. Our results indicated that when incubated without a mediator, the laccases from *P. tephropora* TFRI 707 removed 83.9% of 300 μg/L of TeBT after 7 days incubation. While the laccases obtained from *A. aegerita* and *Pl. geesteranus* removed 73.7% and 70.4% of 300 μg/L of TeBT after 7 days incubation. When incubated in the presence of 0.5 mM ABTS, the laccases obtained from *P. tephropora* TFRI 707, *A. aegerita*, and *Pl. geesteranus* removed 87.4%, 81.7%, and 78.3% of 300 μg/L of TeBT, respectively, after 7 days incubation. Similarly, in the presence of 0.5 mM 4-HAP, VLA, and SIR, the laccases obtained from *P. tephropora* TFRI 707, *A. aegerita*, and *Pl. geesteranus* removed 73.9-88.9% of 300 μg/L of TeBT. The results indicated that the laccases obtained from *P. tephropora* TFRI 707 were better than the other two enzymes in the TeBT removal when used alone without a mediator. Debutylation activities of the laccases obtained from *P. tephropora* TFRI 707, *A. aegerita* and *Pl. geesteranus* were enhanced the most by the presence of 4-HAP, SIR and VLA, respectively. When incubation without a mediator, the laccases obtained from *P. tephropora* TFRI 707, *A. aegerita*, and *Pl. geesteranus* removed 78.0%, 75.5%, and 72.9% of 1000 μg/L of TePhT after 7 days incubation, respectively. In the presence of 0.5 mM ABTS, VLA, 4-HAP, or SIR, the laccases obtained from *P. tephropora* TFRI 707, *A. aegerita*, and *Pl. geesteranus* removed 84.8%–92.0% of 1000 μg/L of TePhT after 7 days incubation. The efficiency of TePhT removal was not different among these enzymes. Thus, addition of mediator enhanced the dephenylation of the laccases obtained from *P. tephropora* TFRI 707, *A. aegerita*, and *Pl. geesteranus*.

Mixed-function oxygenase systems in fish (37) and rat livers (11) have been reported to metabolize TBT through hydroxylation. Another report mentioned that TPhT is resistant to the analogous monooxygenase attack even though it undergoes dephenylation in rat livers (11). Tsang et al. (7) investigated the biodegradation capacity of TBT by using two green microalgae *Chlorella* species and reported that 27% and 41% of the original TBT (100 μg/L) was transformed by these two green microalgae, respectively. DBT was the end degradation product in one culture of *Chlorella* sp. after 14 days incubation. Thus, degradation by laccases were considerably more efficient than those degradation by microorganisms.

Moreover, the use of mediator compounds in an LMS may generate unexpected toxicity (38). By contrast, natural mediators extracted from renewable sources or plants offer environmental and economic benefits (39). In the present study, the TePhT degradation performances of laccases obtained from *P. tephropora* TFRI 707 and *Pl. geesteranus* improved with the addition of a synthetic mediator, such as ABTS or VLA. These results were similar to those of Weng et al. (28). By contrast, the laccases obtained from *A. aegerita* exhibited superior TePhT degradation performance with the addition of a natural mediator, such as 4-HAP. In any case, the natural mediator 4-HAP increased the TeBT degradation performance of the laccases obtained from *P. tephropora* TFRI 707, *A. aegerita*, and *Pl. geesteranus*. Therefore, the laccases obtained from *P. tephropora* TFRI 707, *A. aegerita*, and *Pl. geesteranus* could effectively dealkylate TePhT and TeBT.

During biodegradation of TeBT/TePhT in this study, concentration of TBT/TPhT and DBT/DPhT were low compared to those of MBT/MPhT. Among the butyltin compounds, the most toxic effects were detected with TBT and DBT, they were toxic even at very low concentrations (0.1-1 μM). In contrast, MBT induced lighter cytotoxicity (40). Thus, during degradation of butyltin or phenyltin compounds by fungal laccases, especially the one produced by *P. tephropora* can quickly convert highly toxic tri-substituted (TBT/TPhT) and di-substituted (DBT/DPhT) compounds into low toxicity mono-substituted (MBT/MPhT) compounds, thus the intermediate compounds were not more toxic during degradation by the laccase. This result indicates the potential of using laccases to treat TPhT- or TBT-contaminated wastewater.

The present study indicated that the use of fungal laccase can degrade organotin compounds without a mediator, however, degradation efficiency of organotin compounds can be enhanced when mediator was added. Among the laccases produced by three different kinds of fungi, the laccase of the white rot fungus *P. tephropora* has a better degradation efficiency of organotin compounds. Application of fungal laccases has much better removal efficiency in antimicrobial agent organotin compounds than bacteria or microalgae.

## ACKNOWLEDGEMENTS

The authors thank Professor Ming-Guo Jiang, Guangxi University for the laccase enzyme samples of *A. aegerita* and *Pl. geesteranus*.

